# BPabZIP, a new bZIP protein motif that promotes binding near, and displacement of, nucleosomes

**DOI:** 10.64898/2026.05.20.725981

**Authors:** Desiree Tillo, Victor B. Zhurkin, Aleksey Porollo, Stewart Durell, Hayley K. Hesse, Matthew R. Hass, Phillip J. Dexheimer, Leah C. Kottyan, Matthew T. Weirauch, Charles Vinson

## Abstract

Many transcription factors (TFs) bind only a subset of their canonical binding sites in mammalian cells. To identify differences between bound and unbound sites we examined Zta(N182S), a mutant of the Epstein Barr Virus (EBV)-encoded Zta bZIP protein that binds distinct DNA sequences that are not strongly bound by any known human or viral TF, reducing the effects of selective pressure on endogenous genomic binding sites. We stably expressed Zta(N182S) in human HEK293 cells and monitored protein binding (ChIP-seq) and effects on chromatin accessibility (ATAC-seq). Zta(N182S) binds ~10% of the 14,979 genomic occurrences of the canonical 9-mer ATCACTCAT, creating stronger overall ATAC-seq signal compared to control cells, suggesting nucleosome displacement. Nucleosome occupancy, either predicted or experimentally determined (MNase), indicates that canonical Zta and Zta(N182S) sites are more strongly bound when they are ~60bp from a positioned nucleosome dyad. These data suggest that Zta and Zta(N182S) binding results in nucleosome remodeling, consistent with pioneer-like activity. Examination of amino acids across Zta and human bZIPs identifies four conserved basic amino acids, a proline, and acidic amino acids immediately N-terminal of the basic amino acids of the bZIP domain (PA<u>RR</u>T<u>RK</u>PQQ<u>PE</u>SL<u>EE</u>C<u>D</u>S<u>E</u>L<u>E</u>IKRYKN). We term this new protein motif “BPabZIP” (Basic-Proline-acidic bZIP). Molecular structure predictions for both Zta and human Fos/Jun reveal the basic amino acids interacting with the acidic patch on the nucleosome. The acidic amino acids act as an a-helical extension of the basic region that mimics DNA by interacting with histones H2A and H2B. Taken together, our analyses of this synthetic TF reveal a pioneer-like mechanism that is present in both human and viral bZIP proteins.

## Introduction

Eukaryotic genomes use transcription factors (TFs) that bind specific double stranded DNA (dsDNA) sequences termed transcription factor binding sites (TFBSs) to regulate gene expression (1). In yeast, the bZIP transcription factor GCN4 binds all occurrences of the best-bound 9-mer (2). In mammals, which have ~ 250-fold longer genomes, only a subset of best-bound DNA sequences are occupied by most TFs (3-5).

Several factors are known to contribute to whether a TF binds canonical motifs in mammalian genomes: 1) Mammalian genomes have a bipartite structure (6). There are 20,000 - 50,000 CpG Islands (CGIs) interspersed with vast stretches of A/T rich sequences. CG dinucleotide containing sequences are ~ 10-fold less abundant than sequences without CGs, drastically narrowing the search space for TFs that bind to CG containing k-mers; 2) The CGIs have stable nucleosomes, while the AT rich genome has variably positioned nucleosomes (7, 8). The stable nucleosomes often regulate gene expression by burying or exposing TFBSs (9, 10); 3) TFBSs, particularly CG containing TFBS, colocalize with each other (6, 11, 12), forming promoters and enhancers; 4) TF concentration will affect the proportion of sites bound; 5) for some TFs, posttranslational modifications affect binding. To investigate these contributions to sequence-specific binding, we used a mutant Zta bZIP protein that binds novel dsDNA sites (13) that are not bound by any known endogenous human or viral TF.

Zta is a bZIP domain (14) containing TF encoded in the EBV genome that regulates the switch between the latent and lytic phases of the virus life cycle (15-17). The bZIP domain is a ~60 amino acid long bipartite alpha helix (14). The C-terminus of Zta contains a leucine zipper coiled coil that mediates homo- or hetero-dimerization (14). The N-terminus of the bZIP domain is an a-helical extension of the leucine zipper that binds in the major groove of dsDNA with a binding site of 7-10 nucleotides (14, 18). The function of the N-terminal end of the long a-helical bZIP domain is not well defined. Zta binds (TGA^C^/_G_TCA), the AP-1 motif, also known as the TPA (12-*O-*Tetradecanoylphorbol-13-acetate)-responsive element or TRE. This motif is recognized by multiple human bZIP TFs and is a focal point of gene regulatory circuits (19). Zta also binds sequences containing methylated CG dinucleotides known as CRE sites (TGACGTCA) (20-22). Previously, we used a Protein Binding Microarray (PBM) platform (23) and globally examined Zta binding to dsDNA and dsDNA containing methylated cytosine in CG dinucleotides (24). As part of an exploration of which amino acids in the Zta bZIP domain mediate sequence-specific dsDNA binding to the TRE (T**G**A^C^/_G_T**C**A), we mutated a conserved asparagine (N) to serine (S). Zta(N182S) preferentially binds a new sequence, T**C**A^C^/_G_T**G**A (13), which is not bound by Zta or any known endogenous human TFs.

In this study, we examined the genomic localization (ChIP-seq) and effects on chromatin accessibility (ATAC-seq) of Zta(N182S) and wild type Zta in human cells. Because Zta(N182S) binds a novel DNA motif, we hypothesized that examination of Zta(N182S) binding patterns might give insights into general contributions to sequence specific binding of TFs. For both Zta and Zta(N182S), we observe that a canonical DNA sequence (TRE for Zta, T**C**A^C^/_G_T**G**A for Zta(N182S)) is preferentially located ~60bp upstream of a positioned nucleosome. ATAC-seq data indicate that the nucleosome essential for binding is subsequently displaced. Examination of conservation patterns of amino acids between the Zta and human bZIP domains identifies a sequence we term the BPabZIP motif, which may facilitate interactions with the positioned nucleosome. AlphaFold3 modeling rationalizes the function of these conserved amino acids. Together, these results suggest that a positioned nucleosome is important for Zta binding to the TRE, and that a specific DNA sequence facilitates nucleosome positioning.

## Results

## Zta(N182S) has a distinct binding profile compared to Zta

Full length Zta and Zta(N182S), each tagged with GFP, were stably expressed in human HEK293 cells. This cell line is heavily characterized, with extensive genomic localization data for many proteins due to its inclusion in the ENCODE project (25), and has an available DNA methylome(26). All newly generated ChIP-seq datasets met or exceeded the quality control standards established by the ENCODE consortium (Supplemental Dataset 1). Zta ChIP-seq experiments produced 83,342 peaks and Zta(N182S) produced 72,518 peaks, with 10,983 Zta specific peaks and 22,280 Zta(N182S) specific peaks (Figure 1, Supplemental Dataset 1). Thus, the Zta and Zta(N182S) datasets constitute a high quality and sufficiently distinct *in vivo* binding landscape to perform comparative motif, nucleosome, and accessibility analyses.

**Figure 1.**
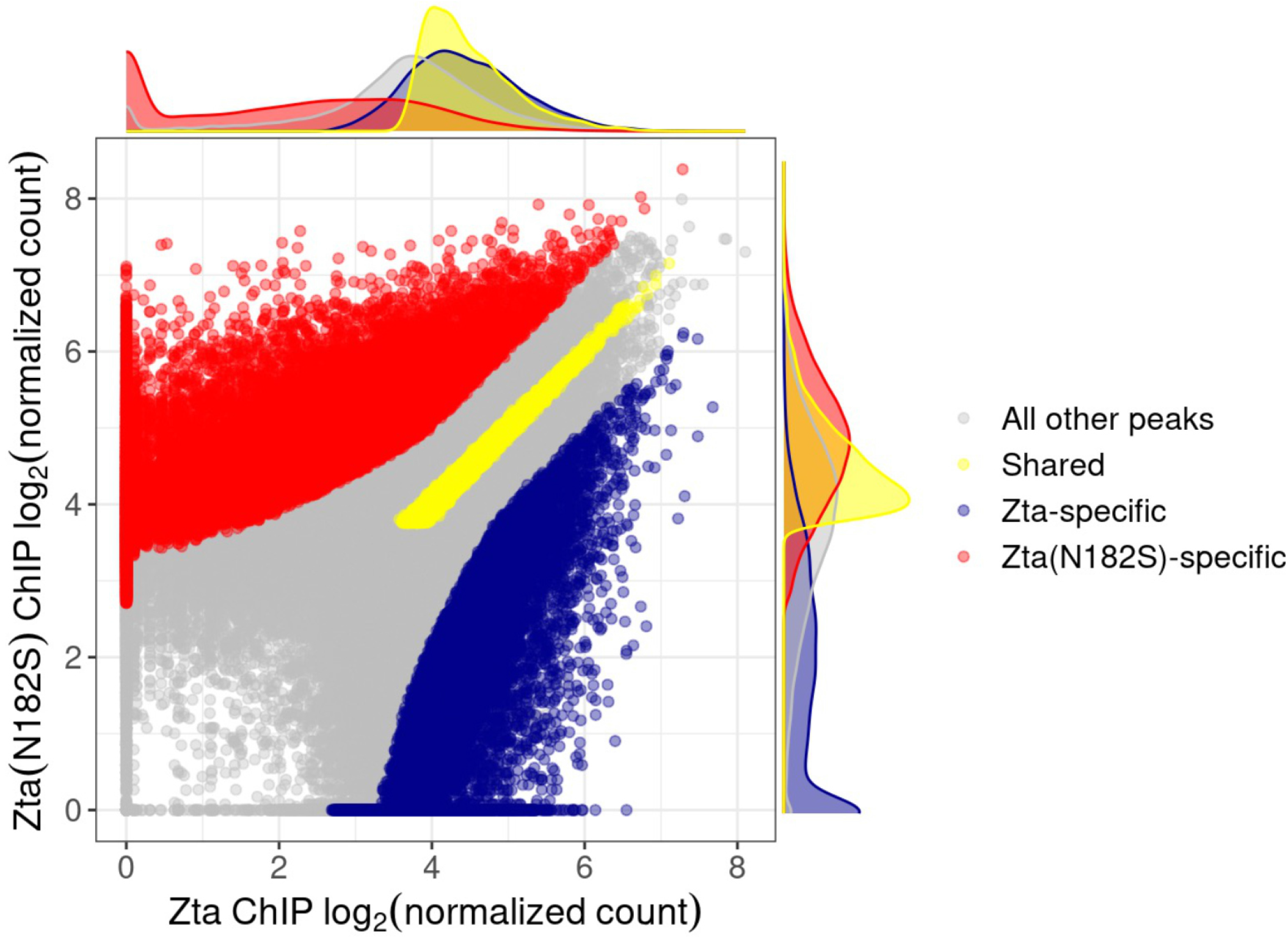
Comparison of Zta(N182S) and Zta ChIP-seq signal in HEK293 cells. Comparison of log2-normalized ChIP-seq signal across a merged set of ChIP-seq peaks for Zta and Zta(N182S). Specific and common peaks are highlighted (See Methods).

### Sequences enriched in Zta and Zta(N182S) ChIP-seq peaks are similar to those measured *in vitro*

We first examined the DNA sequences underlying the genomic positions occupied specifically by either Zta or Zta(N182S). To this end, we employed k-mer enrichment analyses. Importantly, Zta and Zta(N182S) ChIP-seq peaks are enriched for the same 8-mers observed biochemically in protein binding microarray (PBM) experiments for both unmethylated and methylated dsDNA, including established Zta binding sites such as TRE (i.e., the TGA^C^/_G_TCA sequence recognized by AP-1 TFs), meTRE (CGAGTCA), and meZRE2 (TGAGCGA), as well as a modified version of the meZRE2 sequence that is preferentially bound by Zta(N182S) (modmeZRE2, TCAGCGA) (Figure 2, Figure S1) (24, 27).

**Figure 2.**
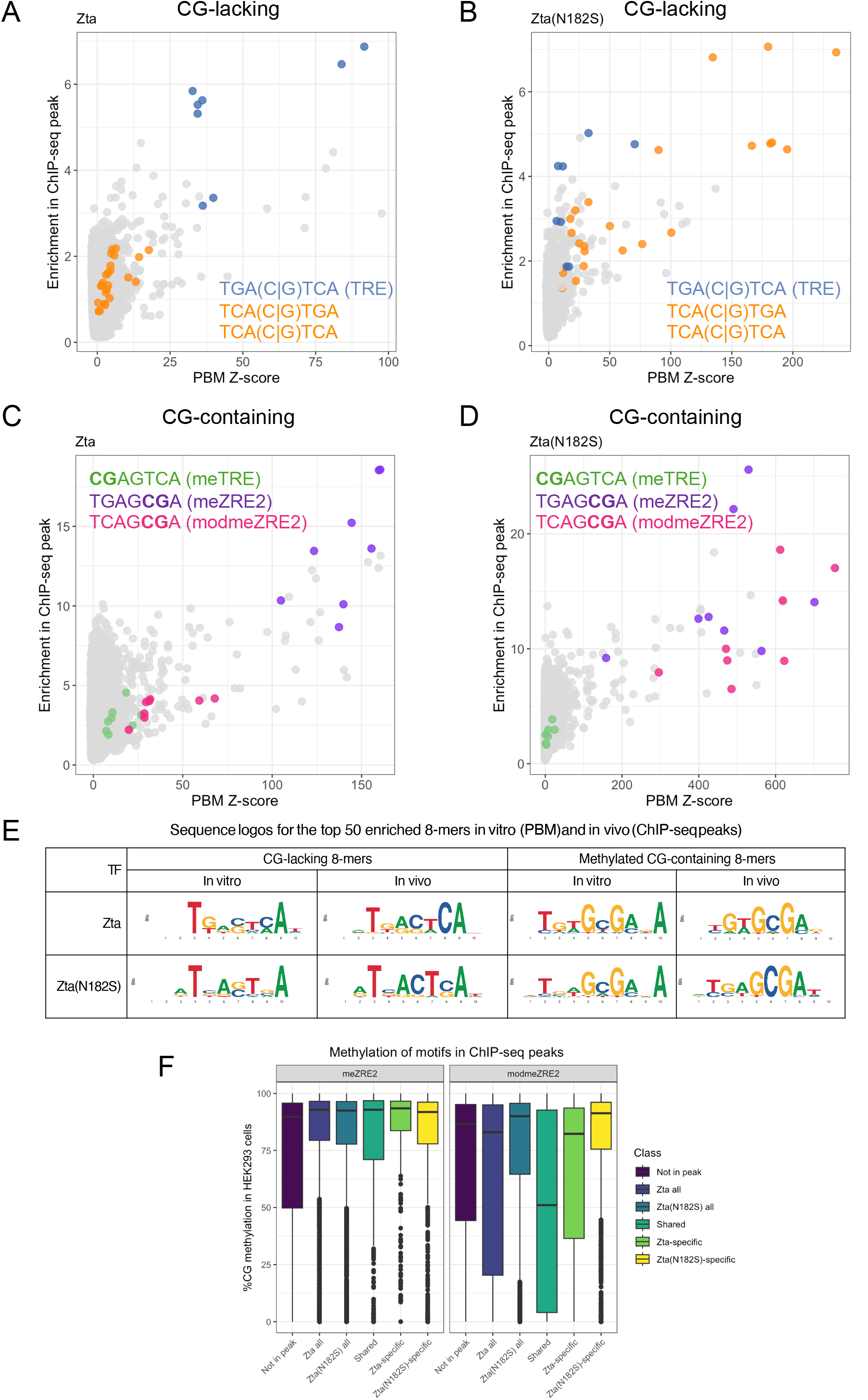
Enrichment of DNA 8-mers in Zta(N182S) and Zta ChIP-seq peaks. (A) Zta binding *in vitro* to 20,349 dsDNA 8-mers lacking a CG dinucleotide in the form of PBM-derived Z-scores (x-axis) (27) vs. enrichment in ChIP-seq peaks (y-axis). (B) Same as in (A) but for Zta(N182S) binding. (C) Zta binding in vitro to 10,762 dsDNA 8-mers with a methylated CG dinucleotide (x-axis) vs. enrichment in ChIP-seq peaks (y-axis). (D) Same as in (C) but for Zta(N182S) binding. (E) Sequence logos generated for the top 50 8-mers bound *in vitro* (PBM) or enriched *in vivo* (ChIP-seq peaks). (F) *In vivo* methylation levels in HEK293 cells (26) of meZRE2 and modmeZRE2-containing 8-mers found in ChIP-seq peaks.

For the 20,349 DNA 8-mers lacking a CG dinucleotide, the most highly enriched 8-mers for Zta include the TRE (AP-1) sequence, as is observed in *in vitro* PBM experiments (Figure 2A, S1A). In contrast, Zta(N182S) enriches for 8-mers containing the 7-mer T**C**A^C^/_G_T**G**A, which is also observed in PBMs (Figure 2B, S1A). For the 10,762 dsDNA 8-mers containing a CG dinucleotide, the most enriched 8-mers in Zta peaks include the methylated meZRE2 (TGAG**CG**A, Figure 2C, S1B) and modmeZRE2 (T**C**AGCGA, Figure 2D, S1B) motifs, as observed biochemically. Sequence logos generated using the top 50 non-CG or CG containing 8-mers recapitulate the *in vitro* results (Figure 2E). Based on publicly available whole genome bisulfite sequencing data (26), CG containing 8-mers that are bound by Zta or Zta(N182S) are indeed methylated in the genome (Figure 2F). In conclusion, the genomic sites occupied by Zta and Zta(N182S) are enriched for the same sequence classes identified biochemically, indicating that the altered binding landscape likely reflects genuine sequence specificity rather than redistribution due to other mechanisms.

### Strongly bound Zta(N182S) motifs show increased ATAC-seq accessibility

We next performed ATAC-seq to examine chromatin accessibility at baseline in HEK293 cells expressing GFP controls or upon expression of either GFP-Zta(N182S) or GFP-Zta (Figure S2, Figure 3). All ATAC-seq datasets meet or exceed the data quality recommendations of the ENCODE consortium (Supplemental Dataset 1). For both proteins, binding to the genome coincided with substantial alterations to human chromatin accessibility (Figure S2).

**Figure 3.**
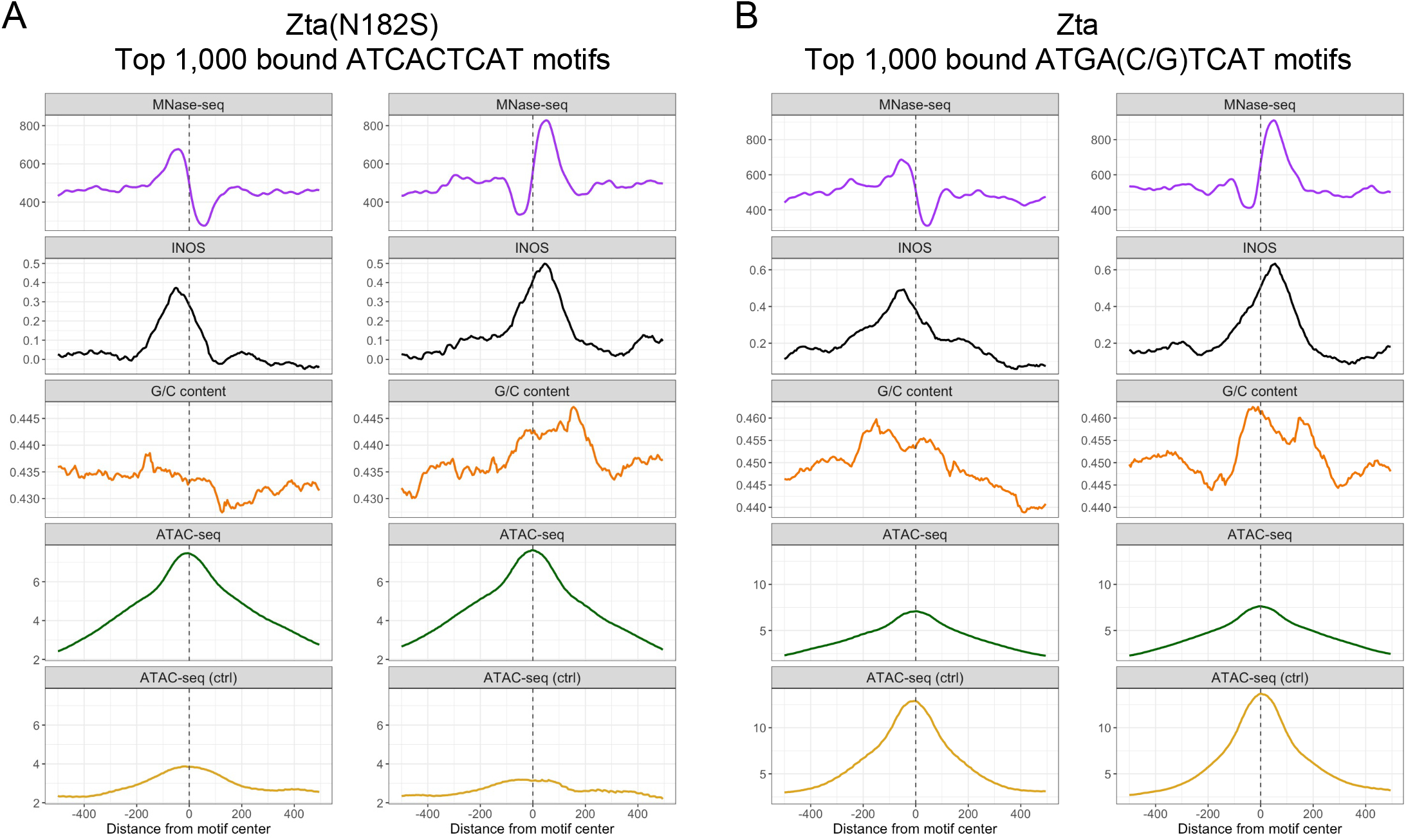
Nucleosome organization surrounding occurrences of the Zta(N182S) and Zta consensus 9-mers. (A) Average MNase signal, INOS, G/C content in 141bp windows, ATAC signal in HEK293 cells expressing Zta(N182S) (ATAC) and control cells (ATAC-ctrl) surrounding the top 1,000 occurrences by Zta(N182S) ChIP-seq signal strength of the 9-mer ATCACTCAT. Occurrences were separated into two groups according to the location of the highest MNase signal (upstream or downstream of the 9mer). (B) Same as in A but for the top 1,000 Zta-bound occurrences of the TRE 9-mer ATGA(C/G)TCAT.

We first examined the ATAC-seq signal in the 1KB region surrounding the 1,000 strongest and the 1,000 weakest bound 9-mers by ChIP-seq signal for Zta(N182S) and Zta (Supplemental Dataset 2, and Figures 3, S3, S4). In cells expressing Zta(N182S), the ATAC-seq signal is strong for the top 1,000 out of 14,979 occurrences in the genome for the 9-mer ATCACTCAT, which is preferentially bound by Zta(N182S) (Figure 3A, S3A). This is also observed for additional strongly bound 9-mers including meZRE2 ATGAG**CG**AT (Figure S5,S6), modmeZRE2 (AT**C**AGC**G**AT, Figure S7), AT**C**AGTCAT (Figure S8) and AT**C**A(C/G)TGAT (Figure S9). These sequences show low accessibility in control cells, suggesting limited pre-existing occupancy at these locations in HEK293 cells, with limited chromatin accessibility in the vicinity of these 9-mers before Zta(N182S) expression (“ATAC-ctrl” in Figure 3A). In contrast, in cells expressing Zta, the strongest bound unmethylated TRE 9-mer (ATGA(C/G)TCA) has strong ATAC-seq signal both in Zta expressing cells and in the control cells (Figure 3B, S4A), suggesting that there is endogenous binding of cellular TFs to this sequence, consistent with the fact that many human TFs such as AP-1 (JUN, FOS, ATF, etc), are known to bind to the TRE (19) (Figure 3B, S4). Taken together, these results support the idea that the strong binding of bZIP proteins to their cognate binding sites correlates with high chromatin accessibility measured by ATAC-seq, suggesting that these factors can impact chromatin organization.

### Zta(N182S) and Zta bind to the edge of nucleosomes

Our analyses show that Zta and Zta(N182S) localize in the genome to the same DNA sequences that are best bound biochemically and that there is chromatin reorganization surrounding these sequences upon Zta(N182S) binding. However, only ~10% of the best bound DNA sequences in the human genome are located in these ChIP-seq peaks (Supplemental Dataset 2). Because only a minority of optimal binding sites are occupied, we next assessed whether local nucleosome architecture might distinguish strongly bound from weakly bound sites. Nucleosome occupancy measured by MNase-seq shows strong MNase signal that covers ~200 bp centered on the preferred 9-mers for both Zta(N182S) and Zta (Figure 3, S3, S4). For the 1,000 weakest bound sites, the MNase signal is less than half that observed for the best bound sites, with no apparent peak near the 9-mers (Figures S3, S4). Similar results were obtained from computing predicted nucleosome occupancy using a DNA-sequence based model (intrinsic nucleosome occupancy scores, INOS, Figures 3, S3, S4) (28), reinforcing that nucleosomes are largely localized according to their intrinsic DNA sequence preferences.

We hypothesized that the high MNase signal and computed INOS score centered on the preferred 9-mers may be due to averaging the nucleosome signal on either side of the 9-mer (i.e., averaging a bimodal). To distinguish a truly centered nucleosome from an average of nucleosomes positioned on either side of the binding site, we oriented each 9-mer by the side with higher local MNase or INOS signal in a 20bp window. The upstream and downstream sides were selected using the top strand of genomic DNA in the reference genome. This approach revealed a dramatically different nucleosome pattern for the strongest-bound 1,000 occurrences for both Zta(N182S) and Zta (Figure 3). For both the experimental MNase data and sequence-based INOS calculations, there is a strong peak of nucleosome occupancy 60-70 bps away from the 9-mer in each direction. Zta and Zta(N182S) binding to any of their preferred 9-mers shows similar patterns of nucleosome occupancy (Figure S3-S9). The 1,000 weakest bound 9-mers show a dramatically different pattern, with low MNase and INOS signal (Figure S3-9). Both lines of evidence support that the weakest bound sites are enriched in regions of the genome that lack localized nucleosomes. GC content, which is a dominant component of the INOS score, does not perfectly correlate with this pattern, suggesting that other components of the INOS score (structural properties, nucleosome disfavoring sequences) contribute. In examining the ATAC-seq signal using oriented alignment, we do not observe asymmetry (Figure 3), suggesting that the localized nucleosome that positively correlated with TF binding is displaced. In conclusion, positioned nucleosomes distinguish strongly bound from weakly bound binding site occurrences, and oriented analysis places the preferred binding sites near nucleosome edges.

### The BPabZIP motif: a conserved basic cluster and acidic helical extension

The ChIP-seq and MNase data indicate that Zta(N182S) preferentially binds near a nucleosome and the ATAC-seq data suggest that the nucleosome is then displaced. We considered whether amino acids immediately N-terminal of the bZIP domain may interact with the nucleosome. To explore this possibility, we examined conservation of this region across Zta and human bZIP proteins. Figure 4A shows a similarity-based phylogenetic tree based on multiple sequence alignment of the 28 amino acids N-terminal of the invariant asparagine (N182) of the Zta bZIP domain across the 54 human bZIP domains. All but four proteins have a set of basic amino acids (K/R) immediately upstream of the asparagine residue (marked as “b”) that is part of the bZIP domain. These K/R residues facilitate interaction of α-helices with DNA phosphates (29) and play a role in nuclear localization (30). Clustering identifies one group of human bZIP domains, which includes FOSB, ATF2 and CREB5, (Group 1, Figure 4A), that contains a series of acidic amino acids (E/D, marked as “a”) immediately N-terminal of the basic region. For this group, another basic cluster (K/R, “B”) lies further toward the N-terminus with an intervening proline (Figure 4A, B). The two motifs frequently occur together, although the distance between them varies. The “B” motif is less defined in the other major cluster identified (Group 2, Figure 4B). In Zta, these sequences are PA<u>RR</u>T<u>RK</u>PQQP**E**SL**EE**CDS**E**L**E**IKRYKN, with the acidic series in bold, and the basic cluster underlined. We designate this the BPabZIP domain. Taken together, this analysis identifies a subset of human bZIP proteins that share a composite N-terminal sequence architecture with Zta, nominating a candidate domain that could contribute to nucleosome-proximal binding.

**Figure 4.**
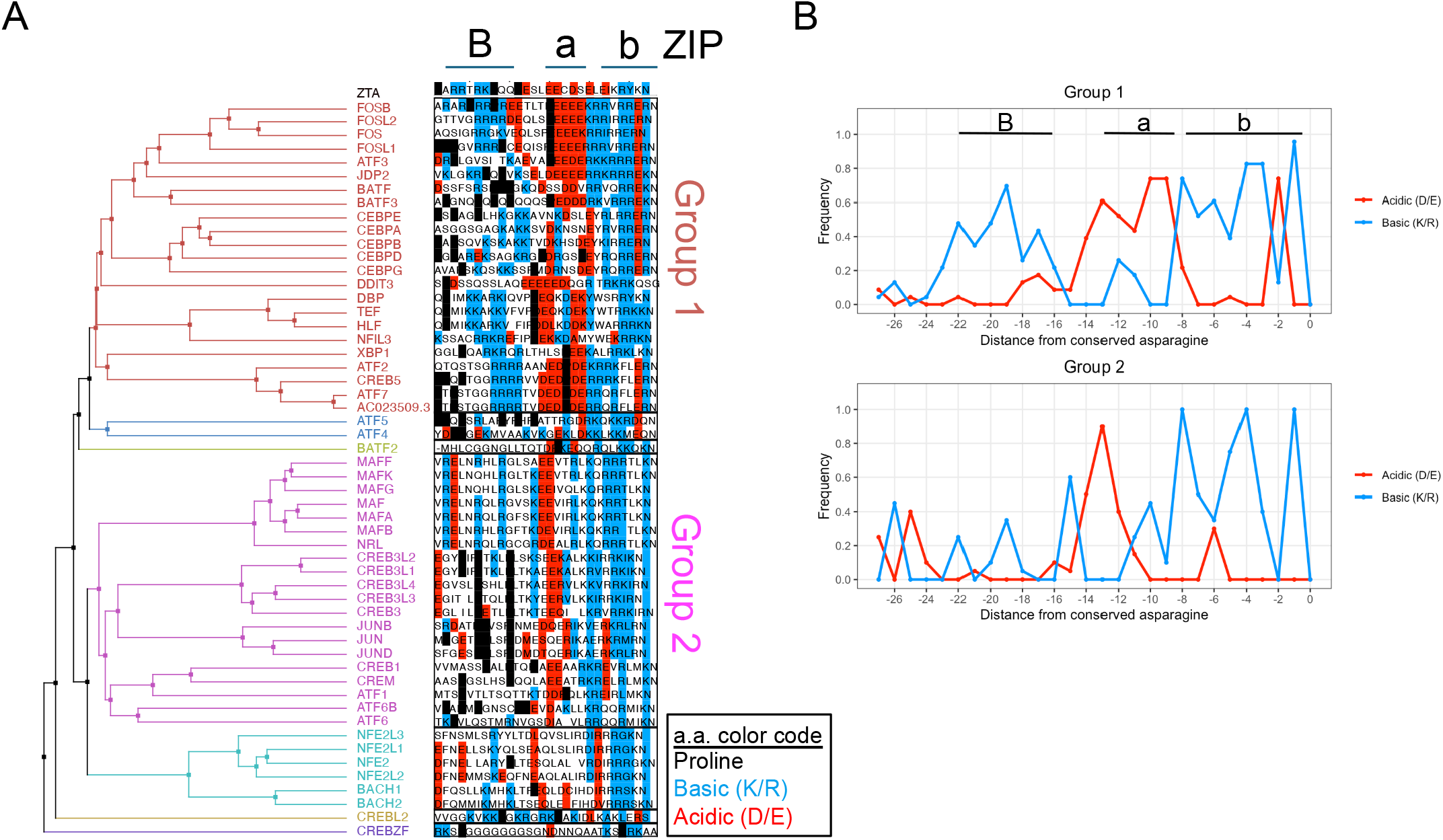
Conservation of the BPa subdomain in human bZIP proteins. (A) The 28 amino acids upstream of the invariant asparagine of the 54 human bZIP proteins and Zta. Acidic amino acids (D/E) are colored in red, basic amino (K/R) acids are colored in blue, and prolines are colored in black. Sequences are clustered according to similarity, and a dendrogram constructed using the BLOSUM62 distance matrix is shown at the left. The approximate location of the acidic extension (“a”) and upstream basic cluster (“B”) are indicated at the top. (B) Frequency of acidic and basic amino acids upstream of the asparagine in the 54 human bZIP proteins in the two largest identified groups.

### Molecular models of the BPabZIP motif of Zta and human FOS/JUN binding a nucleosome

To ask whether the newly identified BPabZIP region could plausibly mediate nucleosome-proximal binding, we used AlphaFold3(31) to generate structural models of Zta BPabZIP segments in the context of a nucleosome and then examined whether similar interactions were also supported for a human FOS/JUN heterodimer. We first examined the interactions between the human nucleosome core particle (histone octamer and 147bp of DNA, Supplemental Dataset 3) and added only the 20 amino acids in the newly identified BPa subdomain of the BPabZIP domain of Zta (Zta20). An electrostatic depiction of Zta20 shows the basic cluster interacting with the acidic patch of the histones H2A/B, a common binding site for many chromatin modifiers (32, 33) (Figure S10). We next generated predicted structural models incorporating the Zta bZIP domain homodimer (BPabZIP) and sequences derived from the chromatosome structure (PDB structure: 4QLC), which includes the nucleosome (the histone octamer with 168bp of DNA) with or without histone H1. We obtained a structure with one a-helix directed toward the nucleosome and covering the nucleosome. The basic region binds DNA at the start of the nucleosome, but does not displace H1, which binds DNA at the entry and exit sites of the nucleosome. We obtained a similar structure in the absence of H1. We suggest that the structure with Zta and H1 bound may be a transition structure and that DNA may compete for DNA binding and displace H1.

These structures do not contain the Zta DNA binding site, suggesting these interactions are driven by protein-protein interactions. To examine the effect of the binding site, we inserted the Zta binding site into 168bp dsDNA in various positions and obtained a variety of structural models ranging from the Zta dimer binding the site and not interacting with the nucleosome, to the Zta dimer binding the site with basic clusters from both monomers near the acidic patch of the nucleosome. In particular, placing the 9-mer TRE sequence 5-bp from the end of the 168-mer (H1168+TRE5, Supplemental Dataset 3), produces a structure very similar to that obtained in the absence of the binding site (Figure 5A,B). Figure S11 shows the bZIP heterodimer FOS/JUN binding the H1168+TRE5 sequence and modeled with human histones rather than the drosophila histones of the EM structure. FOS, which contains the BPabZIP motif (Figure 4A), binds similarly to Zta, while the JUN monomer with only the bZIP domain does not interact with the nucleosome. In conclusion, structural modeling supports the plausibility of a BPabZIP– nucleosome interaction surface, especially involving the nucleosome acidic patch, and extends that hypothesis from Zta to a subset of human bZIP proteins. Taken together, our results suggest that these structures may be potential intermediates in the process of human BPabZIP binding to DNA in the context of a positioned nucleosome, with subsequent displacement of the nucleosome.

**Figure 5.**
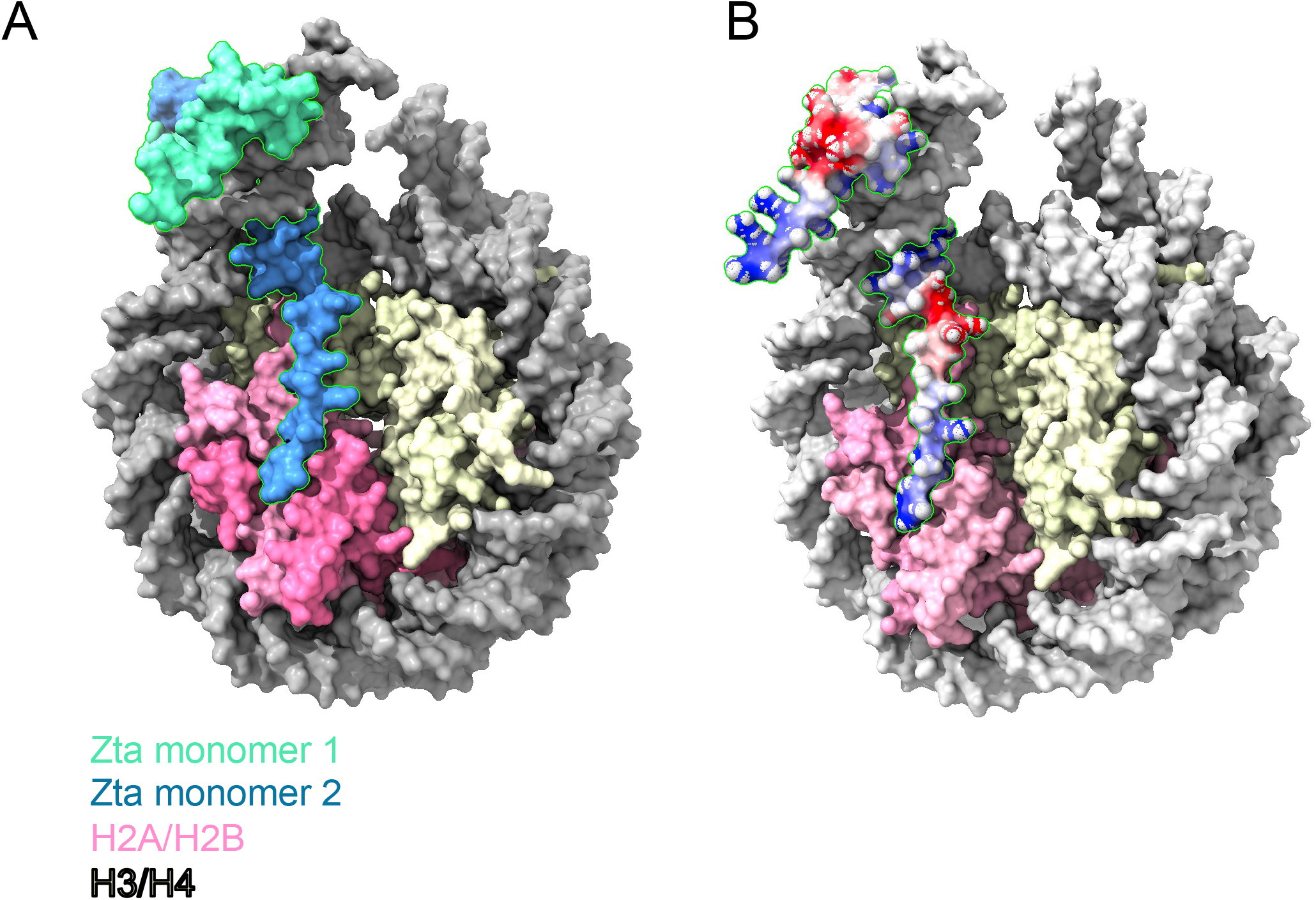
Predicted interactions between the Zta BPabZIP domain and nucleosomes. (A) Molecular structure obtained from Alphafold3 showing each monomer (blue, green) of the Zta BPa subdomain in complex with the nucleosome (4QLC). H2A/B are in pink, H3/H4 are in off-white. (B) Same as in A, with Zta colored according to electrostatic potential, with red, white, and blue denoting negative, neutral, and positive charges, respectively.

## Discussion

In this study, we introduced a foreign DNA-binding transcription factor (Zta(N182S)) into human cells that recognizes DNA sequences not recognized by any known human protein. We examined the relationship between Zta(N182S) binding to canonical 9-mers in human cells and nucleosome position by examining ChIP-, MNase-, and ATAC-seq data. Zta(N182S) has a single amino acid change relative to wildtype Zta that results in new DNA binding specificity (13) and avoids two potential complications in this analysis: (1) Zta(N182S) binds to DNA sequences not bound by any known human TF, avoiding competition with endogenous TF binding; and (2) genomic DNA sequences recognized by Zta(N182S) have not necessarily co-evolved with other TFBSs (11, 34). This system was crucial for revealing that Zta(N182S) binding leads to nucleosome displacement because there is no ATAC-seq signal at the bound canonical Zta(N182S) 9-mers in cells lacking this protein. In contrast, 9-mers bound by Zta, which strongly overlap with human AP-1 TF binding sites, do have an ATAC-seq peak in control cells, indicative of binding by other endogenous AP-1 TFs.

We compared Zta(N182S) binding in cells to nucleosome occupancy measured experimentally (MNase) or determined computationally (INOS) for all occurrences in the genome of several strongly bound 9-mers. Separating occurrences of a specific 9-mer into two groups, those with a higher nucleosome occurrence upstream or downstream of the 9-mer, avoids the signal convolution that may occur if a nucleosome binds either side of the 9-mer and their average produces a peak over the 9-mer by averaging a bimodal (Figure 3, S3-S9). Typically, each non-CG containing 9-mer occurs ~15,000 times in the genome. When we examine the 1,000 best bound occurrences (~7%) of their preferred 9-mer, Zta(N182S) and Zta preferentially bind next to a stable nucleosome. When we examine the 1,000 weakest bound 9-mers, there is no adjacent stable nucleosome. This general pattern is seen for all the 9-mers examined, both unmethylated and methylated.

Zta(N182S) binding creates increased ATAC-seq signal, highlighting that binding near a nucleosome causes nucleosome displacement. In the absence of Zta(N182S), control cells have limited ATAC-seq signal at the best bound 9-mer. After Zta(N182S) expression, there is both a ChIP-seq peak and a strong increase in ATAC-seq signal, indicative of nucleosome displacement. Thus, Zta(N182S) and by extension Zta, have two properties: they bind next to a stable nucleosome and then displace the nucleosome, producing increased chromatin accessibility. For Zta, the best bound TRE 9-mers have increased ATAC-seq signal in control cells, suggestive of endogenous TF binding by AP-1 TFs (19), emphasizing the complexity of examining endogenous bound k-mers.

The term “pioneer factor” was introduced in 2002 (35) for a TF that binds to a nucleosome containing its TFBS (36). In developmental systems, binding of a pioneer factor to a nucleosome is the initial step in a complex rearrangement of chromatin (37). Systematic analysis of TFs binding to DNA wrapped in a nucleosome indicates most classes of TFs do not bind to the DNA on the surface of a nucleosome including most bZIP proteins (38). However, recent work has shown C/EBP, a bZIP protein, binds DNA both at the ends of nucleosomes (39) and over a nucleosome (40). Our analysis suggests that bZIP proteins bind adjacent to nucleosomes *in vivo* and the displacement of the nucleosome suggests that Zta might have pioneer activity in our system.

Examination of conservation patterns between Zta and human bZIP proteins for the 20 amino acids immediately N-terminal of the bZIP domain identified a novel motif that we term BPabZIP. Based on our functional genomics analysis, this motif will be proximal to the nucleosome, which motivated our structural exploration and interpretations. Within this domain, there is conservation of four basic amino acids (B), a proline (P) and an acidic a-helical extension (a) (41) of the basic region (b) of the bZIP domain. Zta(N182S) binding next to a positioned nucleosome could be aided by the basic cluster of BPabZIP binding to the acidic patch on the nucleosome. In this model, nucleosome displacement is achieved by the acidic amino acids (five a-helix forming glutamic acids and one aspartic acid P<u>E</u>SL<u>EE</u>C<u>D</u>S<u>E</u>LUIKRYKN) immediately N-terminus of the basic region, by both binding the histones and repelling the acidic DNA, essentially mimicking DNA (42, 43). A potential function of the intervening proline is to act as an N-terminal cap of the long abZIP helix. This could shorten the distance to the basic cluster and be a force generator helping to unravel the nucleosome. Indeed, our results corroborate a recent study that indicates that amino acid composition rather than the specific sequence of unstructured/disordered domains mediates the pioneer activity of transcription factors (44).

The experimental utility of Zta(N182S) is extensive. Localization to many new locations in the genome enables experiments orthologous to existing regulatory mechanisms. This ability to invade the genome at thousands of locations allows investigators to use their favorite experimental technique or favorite transcription factor to explore this new genome. For example, screens to identify components and factors important in the displacement of the nucleosome can be performed. Other questions include: What additional transactivators regulate gene expression? How can the genome be reorganized? How do you build a regulatory network with specific desired properties? These types of questions can be explored using synthetic proteins such as Zta(N182S).

## Methods

### Plasmid production

A pcDNA5/FRT/TO backbone containing monomeric GFP with a C-terminal linker was used to generate the Zta fusion through TOPO cloning. The N182S mutation was generated through site directed mutagenesis.

### Stable cell generation

Human Flp-In-293 cells were grown on 6 well plates in DMEM supplemented with 10% FBS, 0.1% Anti-anti, and 0.2% normocin. The cells were transfected using Lipofectamine 2000, according to the manufacturer’s protocol, with 0.25 µg of Zta-GFP plasmid and 2.25 mg of the pOG44 Flp-Recombinase expression vector. The media was changed to media containing hygromycin at 50 mg/mL and the hygromycin was gradually increased to 200 mg/mL to produce a pool of cells stably expressing Zta-GFP. The purity of the pool was confirmed by microscopy for GFP. Western blotting using an anti-GFP antibody confirmed that Zta-GFP expression migrated at the expected size.

### ChIP-seq experiments

20 million cells were resuspended in 20 mL of media for a final concentration of 1 million cells/mL and crosslinked with 2 mL of 11% formaldehyde, 0.1M NaCl, 1 mM EDTA, 0.5 mM EGTA, and 50mM HEPES (pH 8.0) while rotating at room temperature for 10 minutes. The crosslinking reaction was stopped through the addition of 1.1 mL of 2.5M glycine and rotating for 5 minutes at room temperature. The cells were washed with cold PBS two times and then pelleted by centrifugation at 5000 RPM for 5 minutes and stored in an Ultra-low freezer until further processing for ChIP. Crosslinked nuclei were isolated by washing the cell pellet with 1 mL of ChIP L1 buffer (10mM HEPES pH 8.0, 140 mM NaCl, 1 mM EDTA, 10% glycerol, 0.25% Triton X-100, and 0.5% NP-40) plus protease inhibitors and rotating at 4 °C for 10 minutes followed by centrifugation at 10,000 g for 5 minutes. Then the nuclear pellets were washed with 1 mL of ChIP L2 buffer (10 mM Tris pH 8.0, 1 mM EDTA, 200 mM NaCl, and 0.5 mM EGTA) plus protease inhibitors by rotating at room temperature for 10 minutes. The nuclei were then pelleted by spinning at 10,000 g for 5 minutes. The nuclei were then washed with 0.5 mL of sonication buffer (TE + 0.1% SDS) with protease inhibitors and again pelleted by centrifugation. The pelleted nuclei were resuspended, followed by resuspension in 1 mL of sonication buffer containing protease inhibitors. The resuspended nuclei were sonicated using a Covaris S220 focused-ultrasonicator at 10% duty cycle, 175 peak power, 200 burst/cycle for 7 minutes. The sonicated chromatin was centrifuged at maximum speed for 10 minutes and the supernatant of soluble chromatin in the supernatant was collected. The salt and detergents in the soluble chromatin were adjusted to final concentrations of 150 mM NaCl, 0.1% NaDOC, 1% Triton X-100, 5% glycerol and protease inhibitors. The chromatin was pre-cleared of Ig by adding 20 μL of Dynabeads Protein A or G and rotating at 4 °C for 40 minutes and centrifuged at 10,000 g for 10 minutes. We performed the ChIP using 200 mL of the pre-cleared chromatin on a SX-8X IP-STAR compact automated system robot from Diagenode with the Abcam anti-GFP antibody along with Dynabeads. We eluted the ChIP material using 10 mM TrisHCl and libraries were prepared by ChIPmentation (45) and sent to the CCHMC Genomics Sequencing Facility for sequencing on an Illumina NovaSeq X Plus (single-end, 100 bp read length).

### ATAC-seq experiments

We followed the OMNI ATAC protocol (46) using 50,000 cells to generate the libraries of open chromatin that were subsequently sequenced. In brief, transposase (Tn5) loaded with sequencing adapter sequences was used to insert sequencing adapters to allow amplification of the accessible DNA, as detailed in (47) Libraries of the accessible DNA sequences were prepared using the OMNI ATAC protocol (46) and sent to the CCHMC Genomics Sequencing Facility for sequencing on an Illumina NovaSeq X Plus (paired-end, 150 bp read length).

### Processing and analysis of ChIP-seq data

ChIP-seq alignment and peak calling was performed using a version of the ENCODE ChIP-seq pipeline to run within a Nextflow environment (48-50). All alignments were performed to the hg38 genome. For all downstream analyses, “overlap optimal” peaks were used. We used the DESEQ2 method contained in Diffbind (51) to determine differentially bound peaks between Zta and Zta(N182S). Peaks were determined to be Zta- or Zta(N182S)-specific if they differed by a two-fold read concentration with FDR<0.01. A random subset of peaks with greater than average read concentration for both Zta and ZtaN182 with <0.1 fold difference was selected as shared peaks (Supplemental Dataset 1).

### Processing and analysis of ATAC-seq data

ATAC-seq alignment and peak calling was performed using a version of the ENCODE ATAC-seq pipeline (48-50) modified to run within a Nextflow environment. All alignments were performed to the hg38 genome. For all downstream analyses, “overlap optimal” peaks were used. For visualization, replicate bam files for all ATAC-seq experiments were merged and converted into signal tracks (bigwig files) normalized to 1X coverage using deeptools v3.5.6 with the following parameters: --binSize 25 --smoothLength 75 --normalizeUsing RPGC --effectiveGenomeSize 2913022398 for hg38.

### Computation of 8-mer enrichment

We filtered out ChIP-seq peaks if >50% of the peak summit (+/-100bp peak summit) was located within repetitive sequence, as defined by RepeatMasker. We counted 8-mers in the repeat-masked and olfactory-gene masked human genome (UCSC build hg38) and those +/-100bp surrounding ChIP-seq peak summits using jellyfish version 2.3.0 (52). We then computed an enrichment score, which is the observed the Observed/Expected ratio of each 8-mer in each peak set (53).

### DNA methylation and MNase-seq data sets

We downloaded whole genome bisulfite sequencing data for HEK293 cells (26) from the Sequence Read Archive (SRA). Raw fastq files were processed using the nf-core/methylseq pipeline v2.6.0 with default parameters. Replicates were merged, and only CpGs covered by at least 10 reads were considered for analysis. Fastq files for MNase-seq in HEK293 cells (38) were downloaded from the European Nucleotide Archive (ENA). Data were processed using the nf-core/mnaseseq pipeline v.1.0.0 with default parameters. Bigwig files representing raw smoothed read density estimates from merged replicates were generated using DANPOS3 software, with the following parameters: --span 1 --smooth_width 20 --width 40.

### Computation of intrinsic nucleosome occupancy score (INOS)

We predicted intrinsic nucleosome occupancy scores (INOSs) at each basepair +/-500 bp of each cognate 9-mer using the Lasso linear model described in Tillo and Hughes(28). For the 500 bp upstream and downstream of each motif occurrence, we calculated the INOS at every base pair using a sliding window of 150 bp.

### Alphafold3 structure predictions

Protein structure predictions were performed using the AlphaFold3 server (31), with the sequences found in Supplemental Dataset 3. 5 predictions were generated per job, with the top structure visualized using ChimeraX software (54) or PyMOL(55).

## Data availability

ATAC-seq and ChIP-seq data have been deposited in the Gene Expression Omnibus (GEO) database under accession codes GSEXXXXXX (Zta) and GSEXXXXXX (Zta(N182S)).

## Acknowledgments

This research was supported in part by the Intramural Research Program of the National Institutes of Health (NIH). The contributions of the NIH author(s) are considered works of the United States Government. The findings and conclusions presented in this paper are those of the author(s) and do not necessarily reflect the views of the NIH or the U.S. Department of Health and Human Services. This work was supported by NIH grants R01 HG010730, R01 AI024717, and R01 NS099068 to M.T.W. and L.C.K. This work utilized the computational resources of the NIH HPC Biowulf cluster (https://hpc.nih.gov).

## Notes

### Competing Interest Statement

The authors have declared no competing interest.

