## Supplemental Figures for "BPabZIP, a new bZIP protein motif that promotes binding near, and displacement of, nucleosomes"

**Supplemental Figure Legends**

**Figure S1. Comparison of enriched 8-mers in Zta and Zta(N182S) ChIP-seq peaks *in vivo*.** (A) Enrichment of all CG dinucleotide-lacking 8-mers in Zta vs Zta(N182S) peaks. (B) Enrichment of all CG dinucleotide-containing 8-mers in Zta vs Zta(N182S) peaks.

**Figure S2. Chromatin accessibility of Zta and Zta(N182S) ChIP-seq peaks.** (A) Heatmap of ATAC-seq signal in Zta- or Zta(N182S)- specific and common peaks. (B) Scatterplot comparing Zta (x-axis) vs. Zta(N182S) (y-axis) normalized ATAC-seq signal in the three classes of peaks as indicated.

**Figure S3**. **Nucleosome organization surrounding occurrences of the Zta(N182S) consensus site ATCACTCAT**. (A) Average MNase signal, INOS, G/C content in 141bp windows, ATAC signal in HEK293 cells expressing Zta(N182S) (ATAC) and control cells (ATAC-ctrl) surrounding the top 1, 000 occurrences by Zta(N182S) ChIP-seq binding signal of the 9-mer ATCACTCAT. (B) Same as in A but separating occurrences into two groups according to the location of the highest MNase signal (upstream or downstream of the 9mer). (C) Same as in B but separating occurrences into two groups according to the location of the highest INOS signal. (D-F) Same as in (A-C) but for the bottom 1,000 occurrences by Zta(N182S) ChIP-seq signal .

**Figure S4**. **Nucleosome organization surrounding occurrences of the TRE 9-mer**. (A) Average MNase signal, INOS, G/C content in 141bp windows, ATAC signal in HEK293 cells expressing Zta (ATAC) and control cells (ATAC-ctrl) surrounding the top 1,000 occurrences by Zta ChIP-seq binding strength of the TRE 9-mer ATGA(C/G)TCAT. (B) Same as in A but separating occurrences into two groups according to the location of the highest MNase signal (upstream or downstream of the 9-mer) (C). Same as in B but separating occurrences into two groups according to the location of the highest INOS signal. (D-F) Same as in (A-C) but for the bottom 1,000 occurrences by Zta ChIP-seq signal.

**Figure S5. Nucleosome organization surrounding occurrences of the meZRE2 9-mer in Zta expressing cells.** (A) Average MNase signal, INOS, G/C content in 141 bp windows, ATAC signal in HEK293 cells expressing Zta (ATAC) and control cells (ATAC-ctrl) surrounding the top 100 occurrences by ChIP-seq binding signal strength of the meZRE2 9-mer ATGAGCGAT. (B) Same as in A but separating occurrences into two groups according to the location of the highest MNase signal (upstream or downstream of the 9mer). (C) Same as in B but separating occurrences into two groups according to the location of the highest INOS signal. (D-F) Same as in (A-C) but for the bottom 100 occurrences by Zta ChIP-seq signal.

**Figure S6. Nucleosome organization surrounding occurrences of the meZRE2 9-mer in Zta(N182S) expressing cells.** (A) Average MNase signal, INOS, G/C content in 141bp windows, ATAC signal in HEK293 cells expressing Zta(N182S) (ATAC) and control cells (ATAC-ctrl) surrounding the top 100 occurrences by ChIP-seq binding signal strength of the 9-mer ATGAGCGAT. (B) Same as in A but separating occurrences into two groups according to the location of the highest MNase signal (upstream or downstream of the site). (C) Same as in B but separating occurrences into two groups according to the location of the highest INOS signal. (D-F) Same as in (A-C) but for the bottom 100 occurrences by Zta(N182S) ChIP-seq signal strength.

**Figure S7. Nucleosome organization surrounding occurrences of the modmeZRE2 9-mer in Zta(N182S) expressing cells.** (A) Average MNase signal, INOS, G/C content in 141bp windows, ATAC signal in HEK293 cells expressing Zta(N182S) (ATAC) and control cells (ATAC-ctrl) surrounding the top 100 occurrences by ChIP-seq binding signal strength of the 9-mer ATCAGCGAT. (B) Same as in A but separating occurrences into two groups according to the location of the highest MNase signal (upstream or downstream of the site). (C) Same as in B but separating occurrences into two groups according to the location of the highest INOS signal. (D-F) Same as in (A-C) but for the bottom 100 occurrences by Zta(N182S) ChIP-seq signal.

**Figure S8. Nucleosome organization surrounding occurrences of ATCAGTCAT in Zta(N182S) expressing cells.** (A) Average MNase signal, INOS, G/C content in 141bp windows, ATAC signal in HEK293 cells expressing Zta(N182S) (ATAC) and control cells (ATAC-ctrl) surrounding the top 1,000 occurrences by ChIP-seq binding signal strength of the 9-mer ATCAGTCAT. (B) Same as in A but separating occurrences into two groups according to the location of the highest MNase signal (upstream or downstream of the site). (C) Same as in B but separating occurrences into two groups according to the location of the highest INOS signal (D-F) Same as in (A-C) but for the bottom 1,000 occurrences by Zta(N182S) ChIP-seq signal.

**Figure S9. Nucleosome organization surrounding occurrences of ATCA(C/G)TGAT in Zta(N182S) expressing cells.** (A) Average MNase signal, INOS, G/C content in 141bp windows, ATAC signal in HEK293 cells expressing Zta(N182S) (ATAC) and control cells (ATAC-ctrl) surrounding the top 100 occurrences by ChIP-seq binding signal strength of the 9-mer ATCA(C/G)TGAT. (B) Same as in A but separating occurrences into two groups according to the location of the highest MNase signal (upstream or downstream of the site). (C) Same as in B but separating occurrences into two groups according to the location of the highest INOS signal. (D-F) Same as in (A-C) but for the bottom 1000 occurrences by Zta(N182S) ChIP-seq signal.

**Figure S10. Predicted interactions between the Zta20 BPab domain and nucleosomes.** (A) AlphaFold3 generated 3D model of the 20 residue BPa subdomain of the Zta BPabZIP domain in interaction with human histone. The 20-residue Zta motif is displayed in balls-n-sticks representation, with positively charged (basic) residues colored light blue and negatively charged (acidic) residues colored pink. The human histone proteins are rendered as molecular surfaces, with basic and acidic residues colored dark blue and red, respectively. The DNA double helix is also displayed in surface representation, with the two strands colored light gray and dark gray. (B) A zoom-in visualization of the same protein complex with the focus on the BPa subdomain.

**Figure S11. Predicted interactions between JUN/FOS and nucleosomes.** AlphaFold3 generated 3D model of the cJun/cFos heterodimer and human nucleosomes.

### Figure S1

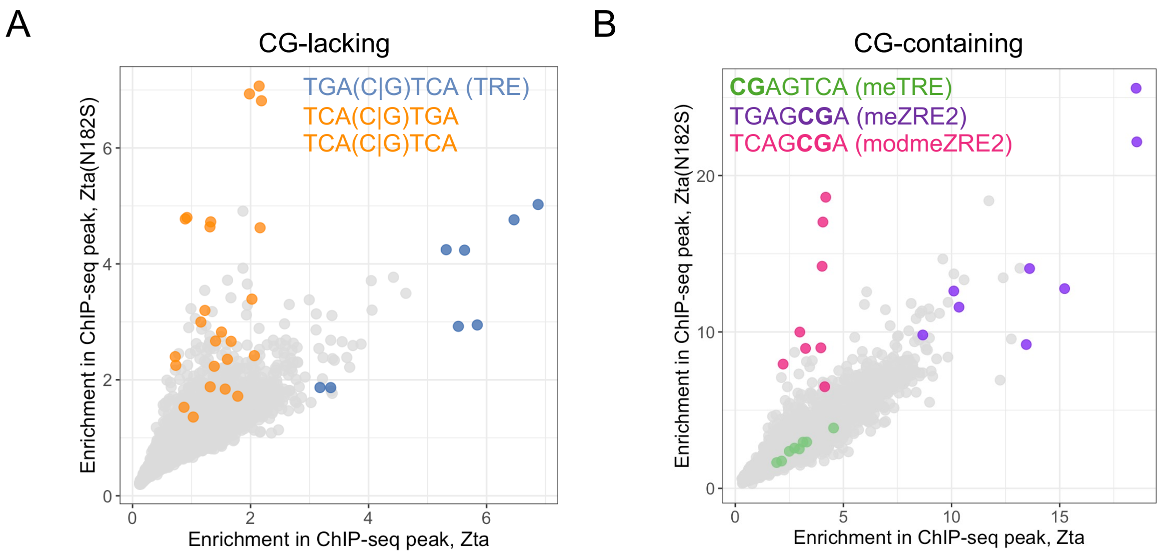
A
CG-lacking
TGA(C|G)TCA (TRE)
TCA(C|G)TGA
TCA(C|G)TCA
Enrichment in ChIP-seq peak, Zta(N182S)
Enrichment in ChIP-seq peak, Zta
0
2
4
6
B
CG-containing
CGAGTCA (meTRE)
TGAGCGA (meZRE2)
TCAGCGA (modmeZRE2)
Enrichment in ChIP-seq peak, Zta(N182S)
Enrichment in ChIP-seq peak, Zta
0
5
10
15
20

Figure S2

A

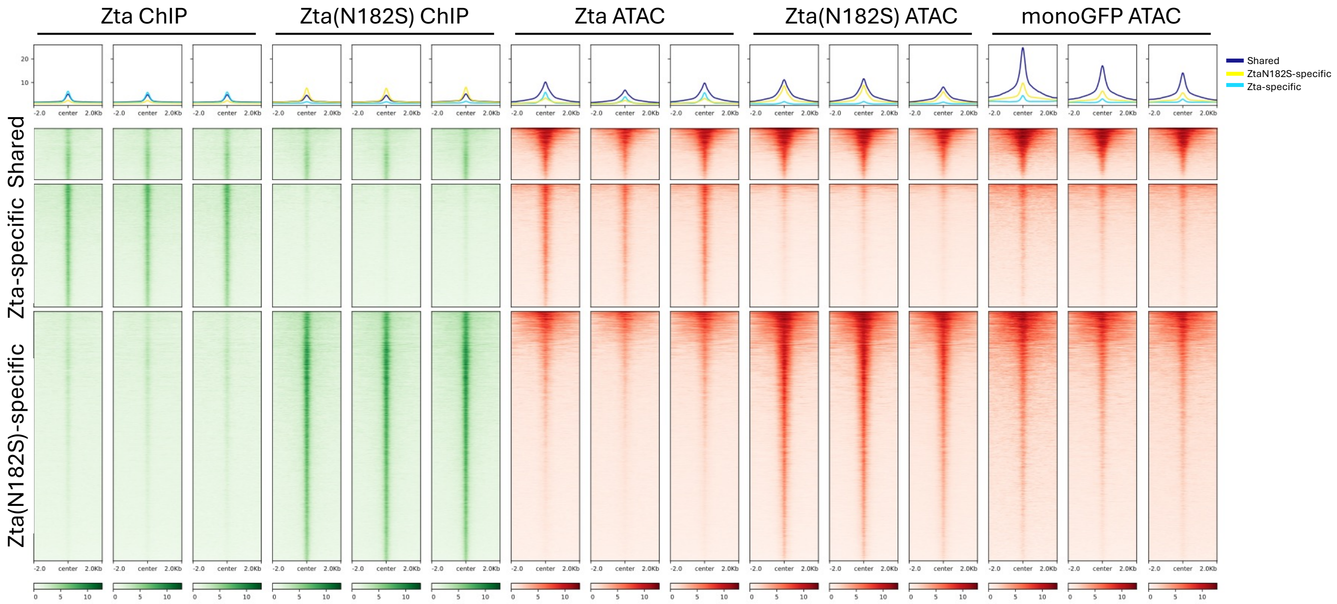
Zta ChIP
Zta(N182S) ChIP
Zta ATAC
Zta(N182S) ATAC
monoGFP ATAC
Shared
ZtaN182S-specific
Zta-specific
Shared
Zta-specific
Zta(N182S)-specific
-2.0
center
2.0Kb

B

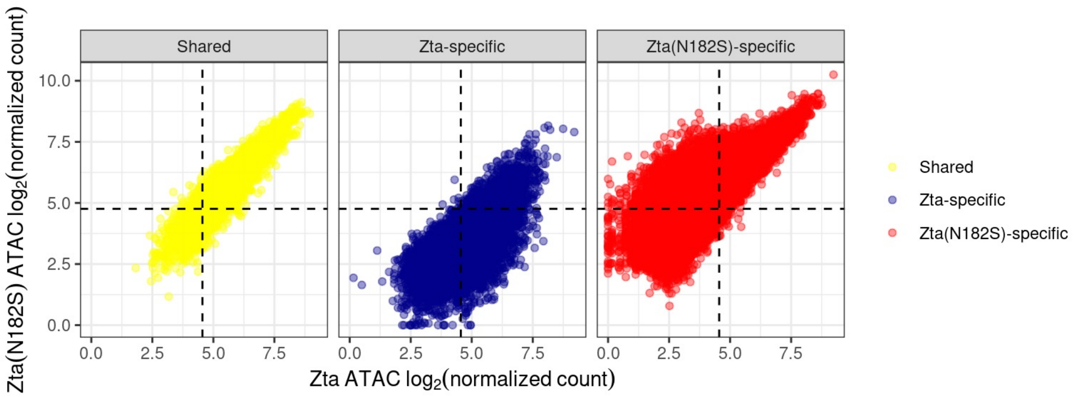
Shared
Zta-specific
Zta(N182S)-specific
Zta(N182S) ATAC log2(normalized count)
Zta ATAC log2(normalized count)
Shared
Zta-specific
Zta(N182S)-specific

### Figure S3

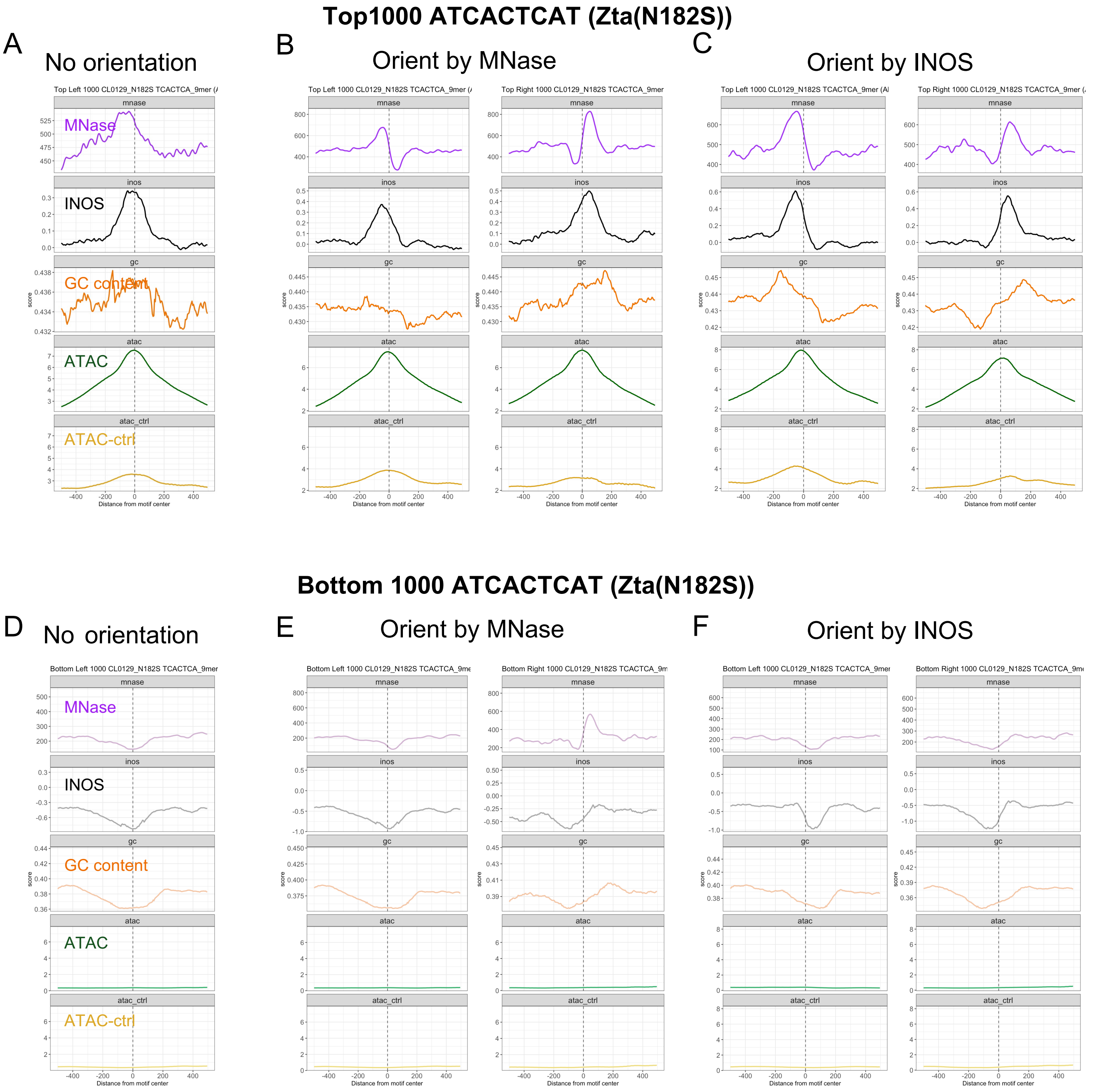

Top1000 ATCACTCAT (Zta(N182S))
A
No orientation
B
Orient by MNase
C
Orient by INOS
MNase
INOS
GC content
ATAC
ATAC-ctrl
mnase
inos
gc
atac
atac_ctrl
score
Distance from motif center
Bottom 1000 ATCACTCAT (Zta(N182S))
D
No orientation
E
Orient by MNase
F
Orient by INOS
MNase
INOS
GC content
ATAC
ATAC-ctrl

Figure S4

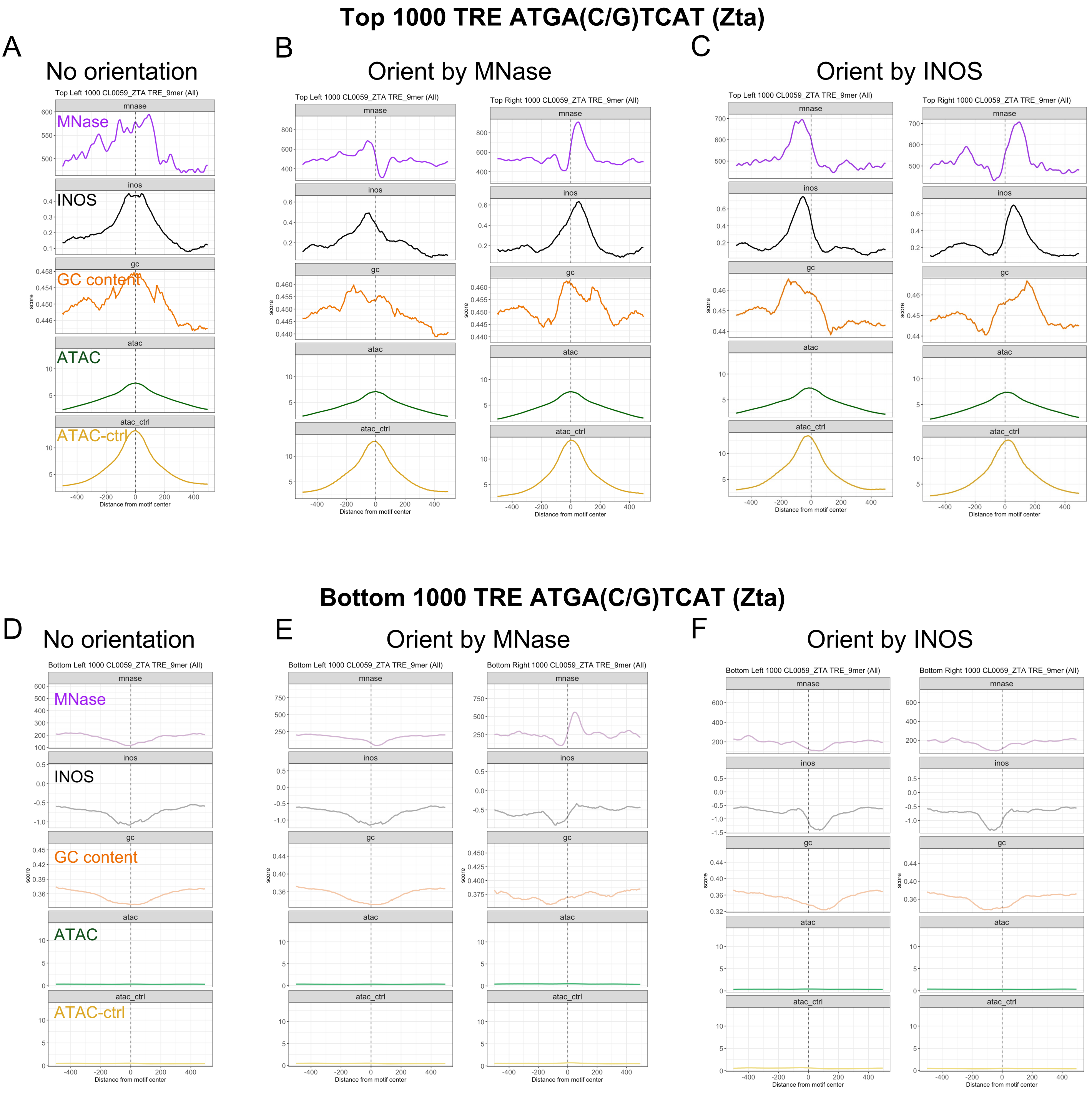

Top 1000 TRE ATGA(C/G)TCAT (Zta)
A
No orientation
B
Orient by MNase
C
Orient by INOS
Top Left 1000 CL0059_ZTA TRE_9mer (All)
Top Right 1000 CL0059_ZTA TRE_9mer (All)
mnase
inos
gc
atac
atac_ctrl
MNase
INOS
GC content
ATAC
ATAC-ctrl
score
Distance from motif center
Bottom 1000 TRE ATGA(C/G)TCAT (Zta)
D
No orientation
E
Orient by MNase
F
Orient by INOS
Bottom Left 1000 CL0059_ZTA TRE_9mer (All)
Bottom Right 1000 CL0059_ZTA TRE_9mer (All)

### Figure S5

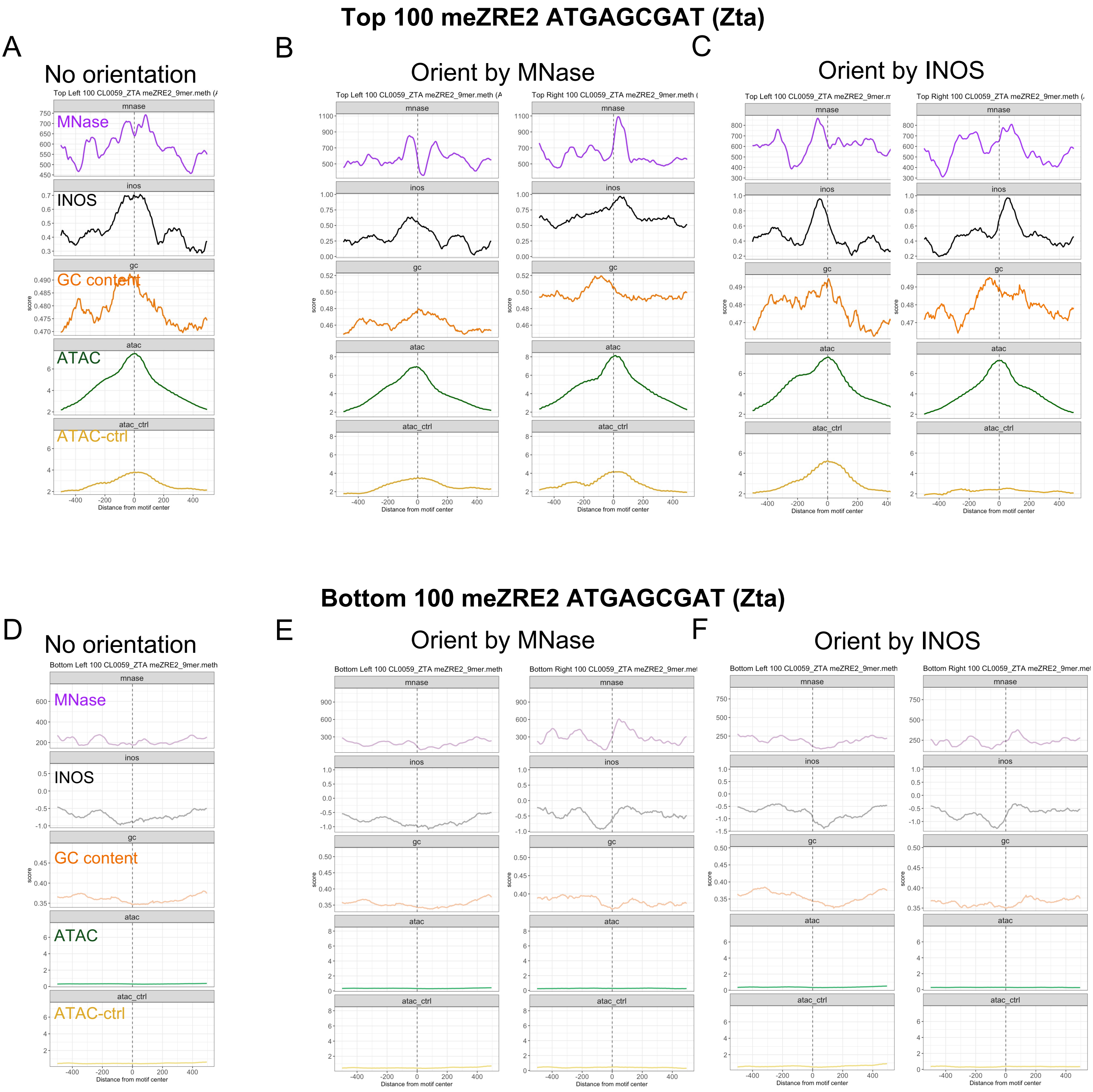
Top 100 meZRE2 ATGAGCGAT (Zta)
A
No orientation
B
Orient by MNase
C
Orient by INOS
MNase
INOS
GC content
ATAC
ATAC-ctrl
mnase
inos
gc
atac
atac_ctrl
score
Distance from motif center
Top Left 100 CL0059_ZTA meZRE2_9mer.meth
Top Right 100 CL0059_ZTA meZRE2_9mer.meth
Bottom 100 meZRE2 ATGAGCGAT (Zta)
D
No orientation
E
Orient by MNase
F
Orient by INOS
Bottom Left 100 CL0059_ZTA meZRE2_9mer.meth
Bottom Right 100 CL0059_ZTA meZRE2_9mer.met

### Figure S6

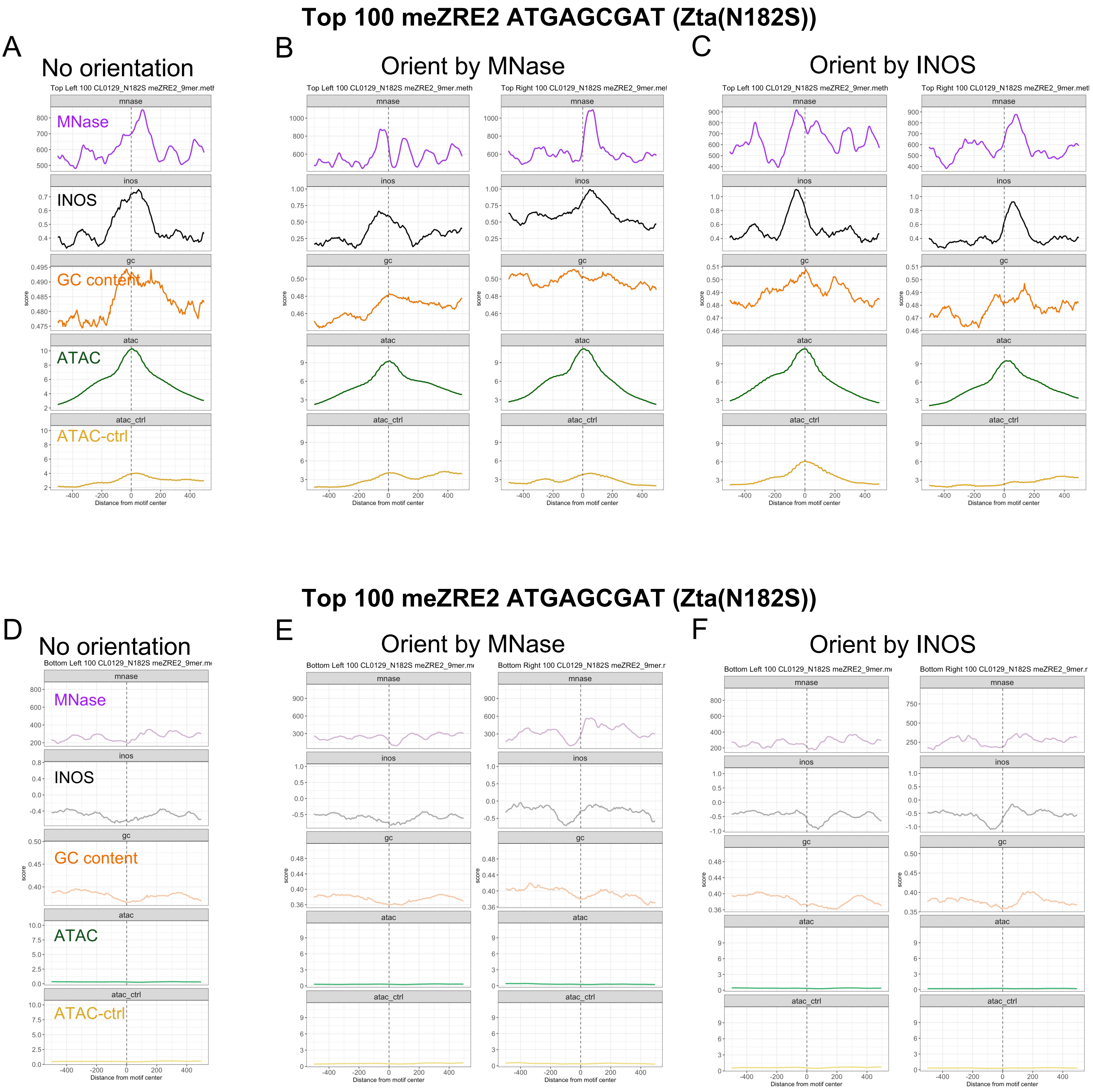
Top 100 meZRE2 ATGAGCGAT (Zta(N182S))
A
No orientation
Top Left 100 CL0129_N182S meZRE2_9mer.meth
B
Orient by MNase
Top Left 100 CL0129_N182S meZRE2_9mer.meth
Top Right 100 CL0129_N182S meZRE2_9mer.met
C
Orient by INOS
Top Left 100 CL0129_N182S meZRE2_9mer.meth
Top Right 100 CL0129_N182S meZRE2_9mer.met
mnase
inos
gc
atac
atac_ctrl
MNase
INOS
GC content
ATAC
ATAC-ctrl
score
Distance from motif center
Top 100 meZRE2 ATGAGCGAT (Zta(N182S))
D
No orientation
Bottom Left 100 CL0129_N182S meZRE2_9mer.m
E
Orient by MNase
Bottom Left 100 CL0129_N182S meZRE2_9mer.m
Bottom Right 100 CL0129_N182S meZRE2_9mer.r
F
Orient by INOS
Bottom Left 100 CL0129_N182S meZRE2_9mer.m
Bottom Right 100 CL0129_N182S meZRE2_9mer.r

### Figure S7

**Top 100 modmeZRE2 ATCAGCGAT (Zta(N182S))**

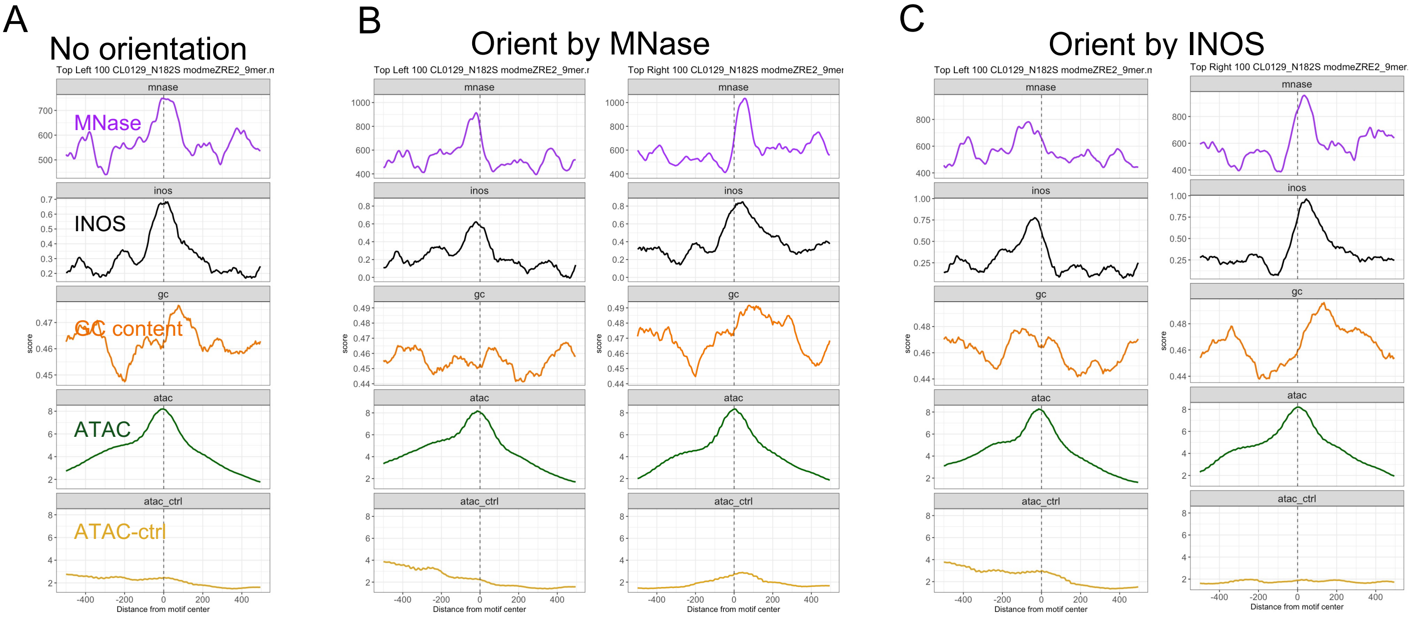


**Top 100 modmeZRE2 ATCAGCGAT (Zta(N182S))**

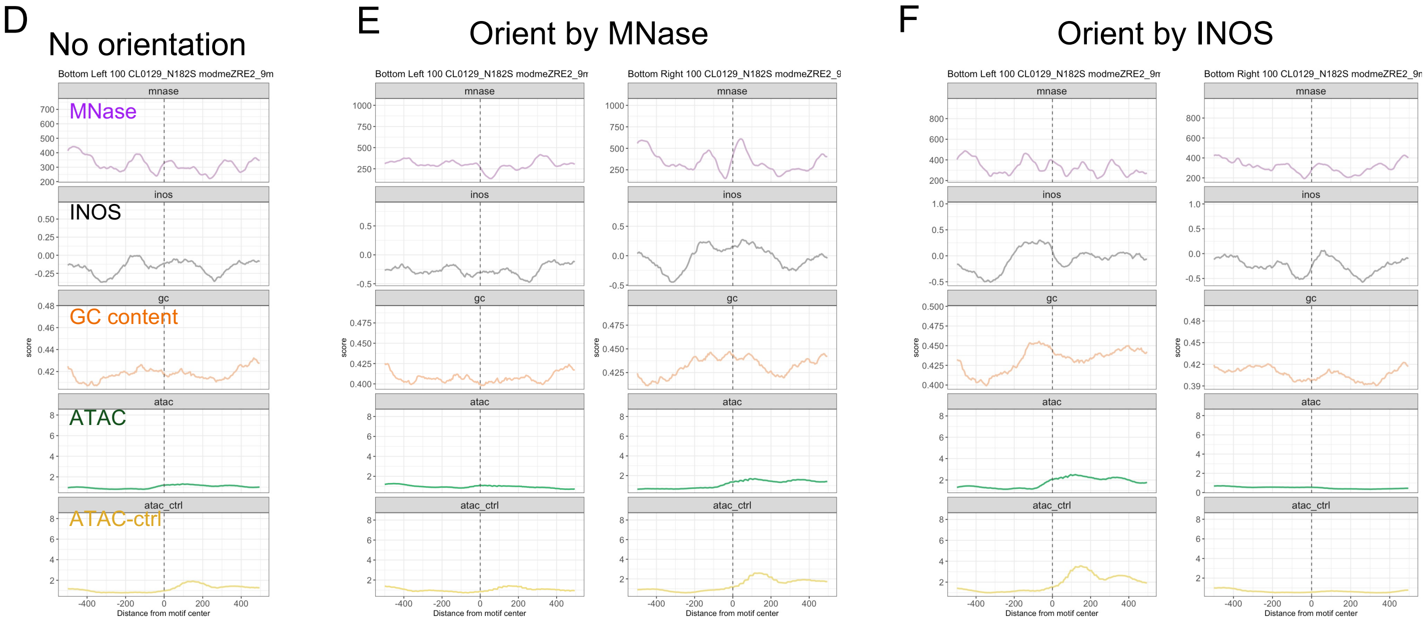

### Figure S8

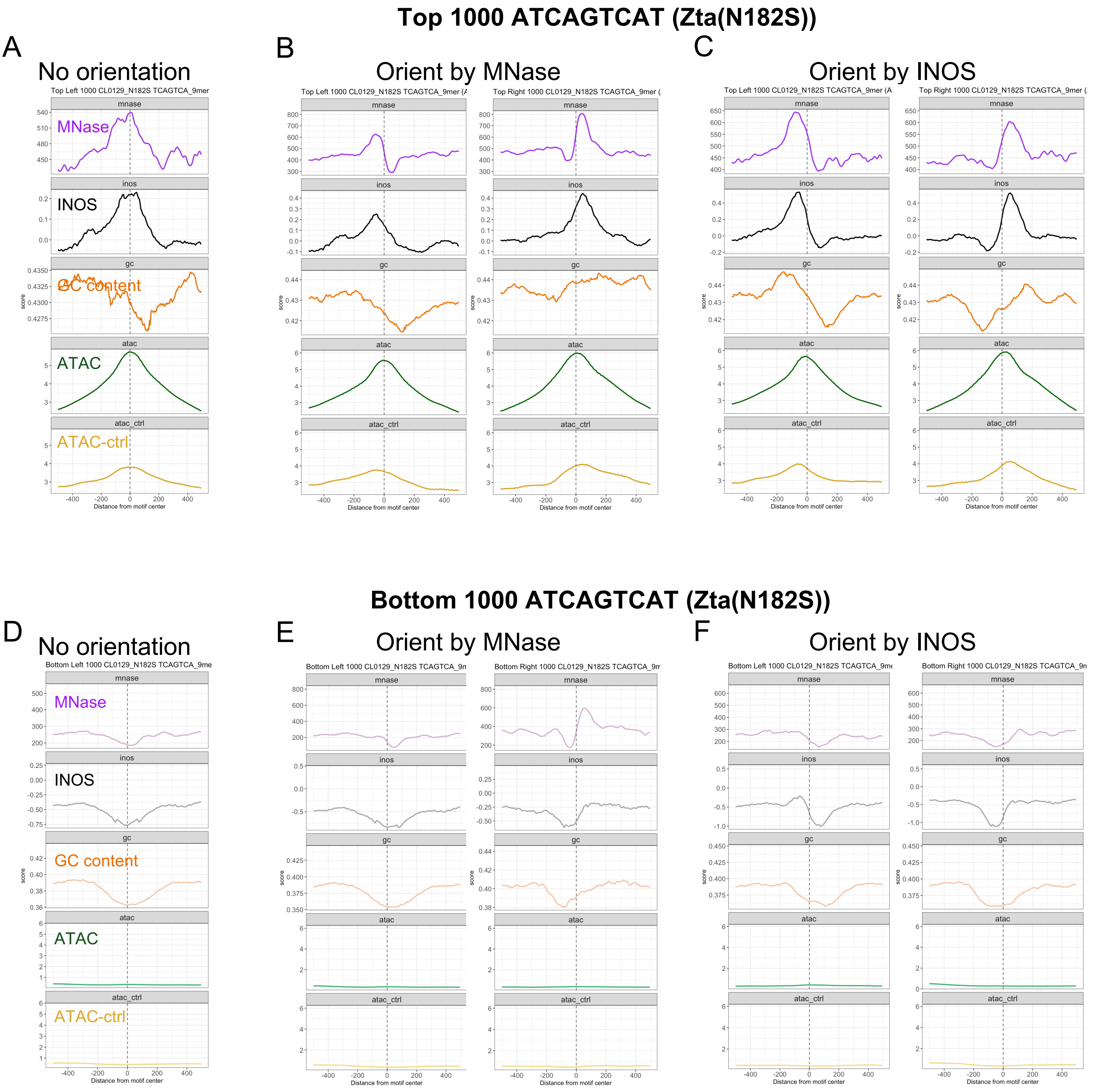

Top 1000 ATCAGTCAT (Zta(N182S))
A
No orientation
Top Left 1000 CL0129_N182S TCAGTCA_9mer
MNase
INOS
GC content
ATAC
ATAC-ctrl
B
Orient by MNase
C
Orient by INOS
mnase
inos
gc
atac
atac_ctrl
score
Distance from motif center
Bottom 1000 ATCAGTCAT (Zta(N182S))
D
No orientation
E
Orient by MNase
F
Orient by INOS

### Figure S9

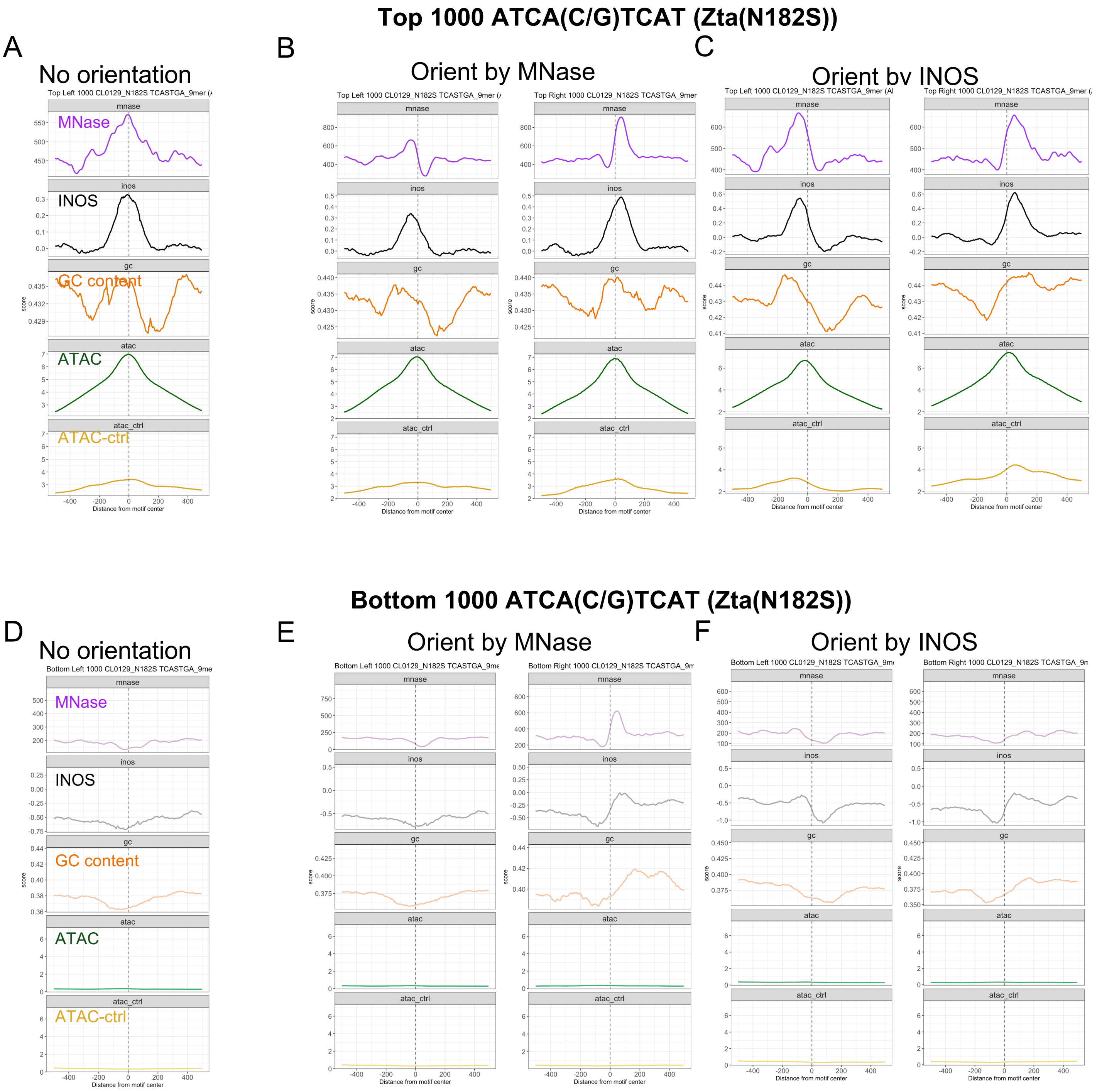

Top 1000 ATCA(C/G)TCAT (Zta(N182S))
A
No orientation
B
Orient by MNase
C
Orient by INOS
Top Left 1000 CL0129_N182S TCASTGA_9mer (A
Top Right 1000 CL0129_N182S TCASTGA_9mer
mnase
inos
gc
atac
atac_ctrl
MNase
INOS
GC content
ATAC
ATAC-ctrl
score
Distance from motif center
Bottom 1000 ATCA(C/G)TCAT (Zta(N182S))
D
No orientation
E
Orient by MNase
F
Orient by INOS
Bottom Left 1000 CL0129_N182S TCASTGA_9me
Bottom Right 1000 CL0129_N182S TCASTGA_9m

Figure S10

A

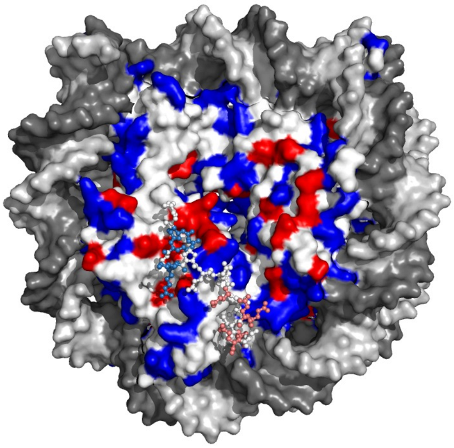

B

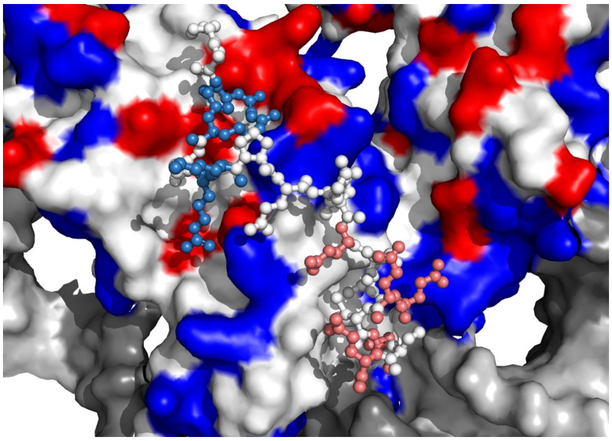

Figure S11

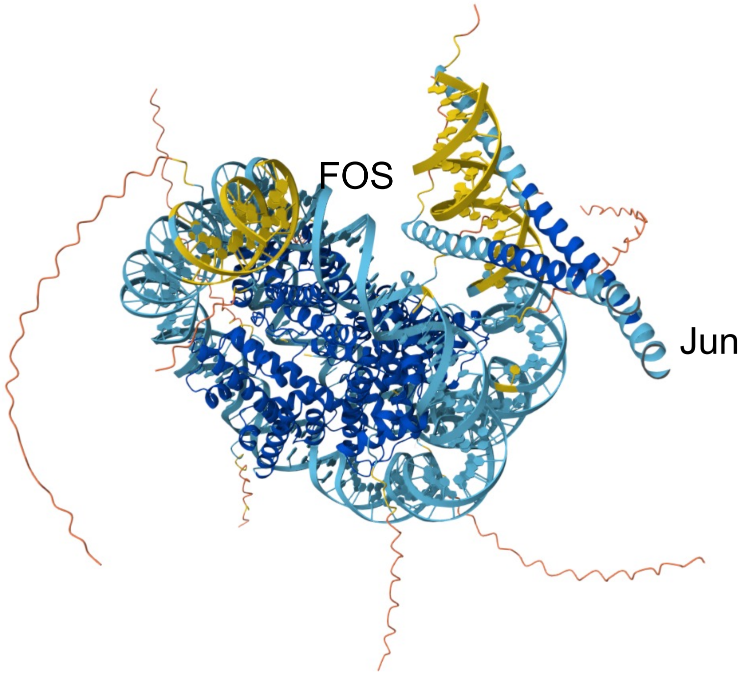
